# Anaerobic growth of *Listeria monocytogenes* on rhamnose is stimulated by Vitamin B12 and bacterial microcompartment dependent 1,2-propanediol utilization

**DOI:** 10.1101/2021.04.20.440696

**Authors:** Zhe Zeng, Siming Li, Sjef Boeren, Eddy J. Smid, Richard A. Notebaart, Tjakko Abee

**Affiliations:** Food Microbiology, Wageningen University and Research, Wageningen, The Netherlands; Laboratory of Biochemistry, Wageningen University and Research, Wageningen, The Netherlands

## Abstract

The food-borne pathogen *Listeria monocytogenes* is able to form proteinaceous organelles called bacterial microcompartments (BMCs) that optimize the utilization of substrates, such as 1,2-propanediol, and confer an anaerobic growth advantage. Rhamnose is a deoxyhexose sugar abundant in a range of environments including the human intestine, and can be degraded in anaerobic conditions into 1,2-propanediol, next to acetate and lactate. Rhamnose-derived 1,2-propanediol has been found to link with BMCs in a limited number of commensal human colonic species and some human pathogens such as *Salmonella enterica*, but the involvement of BMCs in rhamnose metabolism and potential physiological effects on *L. monocytogenes* are still unknown. In this study, we firstly test the effect of rhamnose uptake and utilization on anaerobic growth of *L. monocytogenes* EGDe without and with added vitamin B12, followed by metabolic analysis. We unveil that the vitamin B12-dependent activation of *pdu* stimulates metabolism and anaerobic growth of *L. monocytogenes* EGDe on rhamnose via 1,2-propanediol degradation into 1-propanol and propionate. Transmission electron microscopy of *pdu*-induced cells shows that BMCs are formed and additional proteomics experiments confirm expression of *pdu* BMC shell proteins and enzymes. Finally, we discuss physiological effects and energy efficiency of *L. monocytogenes pdu* BMC-driven anaerobic rhamnose metabolism and impact on competitive fitness in environments such as the human intestine.

## 1. Introduction

*Listeria monocytogenes* is a Gram-positive facultative anaerobe and a food-borne pathogen which causes a severe human infection called listeriosis [1, 2]. The pathogen continues to cause food-borne illness outbreaks characterised by high mortality ranging from 20 to 30% [1, 3]. *L. monocytogenes* is found ubiquitously in natural environments and it can survive a variety of stress conditions leading to the colonization of different niches including a range of food processing environments [1, 3, 4]. To survive in such a variety of niches, *L. monocytogenes* should be able to adapt to environmental stresses and to use a range of nutrients for growth in aerobic and anaerobic conditions [1, 5, 6].

Recent studies on anaerobic growth of *L. monocytogenes* have provided evidence that it has the capacity to form proteinaceous organelles so-called bacterial microcompartments (BMCs) that enable extension of its metabolic repertoire by supporting the utilization of 1,2-propanediol and ethanolamine [7–9]. BMCs are self-assembling organelles that consist of an enzymatic core that is encapsulated by a semi-permeable protein shell [7, 10, 11]. The separation of the encapsulated enzymes from the cytosol is thought to protect the cell from toxic metabolic intermediates such as aldehydes, and prevent unwanted side reactions [7, 10, 11]. In our previous studies, we showed that the *L. monocytogenes* 1,2-propanediol utilization gene cluster (*pdu*) is activated in the presence of 1,2-propanediol and vitamin B12, resulting in stimulation of growth in anaerobic conditions [8]. Vitamin B12 is required for activation of the *pdu* cluster in *L. monocytogenes* [8, 12] and to act as a cofactor of 1,2-propoanediol reductase [13]. Activation of BMC-dependent *pdu* supports degradation of 1,2-propanediol via the toxic intermediate propionaldehyde into 1-propanol and propionate via respective reductive and oxidative branches, with the latter resulting in extra ATP generation leading to enhanced anaerobic growth of *L. monocytogenes* [8]. Notably, 1,2-propanediol is a major end product from the anaerobic degradation of mucus-derived rhamnose by human intestinal microbiota and it is thought to be an important energy source supporting intestinal growth of selected pathogens such as *Salmonella spp*. and *L. monocytogenes* [7, 14–16].

Rhamnose is a naturally occurring deoxyhexose sugar abundant in glycans on surfaces of mammalian and bacterial cells and in cell walls of many plant and insect species [14, 17]. Anaerobic metabolism of rhamnose has been studied previously in a range of bacteria including *E. coli*, and rhamnose is parallelly metabolized into lactaldehyde and dihydroxyacetone phosphate (DHAP) [18, 19]. DHAP is converted in the glycolytic pathway leading to a variety of fermentation products, while lactaldehyde is converted to 1,2-propanediol that is subsequently secreted [18, 19]. Notably, for example in *Salmonella spp*. and *Clostridium phytofermentans*, rhamnose-derived 1,2-propanediol can be converted to 1-propanol and propionate via BMC-dependent *pdu* [14, 16]. Although rhamnose-derived 1,2-propanediol was found to be metabolised via a *pduD*-dependent pathway in *Listeria innocua* [20], the possible activation and contribution of BMC-dependent pdu to anaerobic metabolism and growth of *L. monocytogenes* on rhamnose remains to be investigated.

In this study, we firstly quantified the effect of rhamnose as sole carbon source on anaerobic growth and metabolism of *L. monocytogenes* in absence and presence of vitamin B12 (cobalamine), an essential co-factor of 1,2-propoanediol reductase, the signature enzyme of BMC-dependent pdu [13] Next, we analysed rhamnose utilization and end product formation, and combined with Transmission Electron Microscopy and proteomics, we provide evidence for a B12-dependent *pdu*-induced metabolic shift. We summarize our findings in a model integrating BMC-dependent *pdu* with rhamnose metabolism, and discuss impact on growth and survival of *L. monocytogenes* in anaerobic environments such as the human intestine.

## 2. Materials and Methods

### 2.1 Strains, Culture Conditions, and Growth Measurements

All experiments in this study were carried out with *L. monocytogenes* EGDe anaerobically grown at 30°C in defined medium MWB [21]. Overnight grown cells in Luria Broth (LB) were washed three times in PBS before inoculation into MWB. MWB was supplemented with 20mM L-rhamnose as sole carbon source with or without addition of 20nM vitamin B12. Anaerobic conditions were achieved by Anoxomat Anaerobic Culture System with a gas mixture composed of 10% CO_2_, 5% H_2_, 85% N_2_. MWB with 20 mM rhamnose and 20 nM vitamin B12 was defined as rhamnose *pdu*-induced, while MWB with 20 mM rhamnose was defined as rhamnose *pdu* noninduced condition. OD_600_ measurements in MWB were performed every 12 h for 3 days. Plate counting in MWB to quantity Colony Forming Units (CFUs) was performed every 24 h for 3 days.

### 2.2 Analysis of metabolites for Rhamnose metabolism using High Pressure Liquid Chromatography (HPLC)

Samples were taken from the cultures at 0, 24, 48, and 72 h. After centrifugation, the supernatant was collected for the HPLC measurements of rhamnose, acetate, lactate, 1,2-propanediol, 1-propanol and propionate. The experiment was performed with three biological replicates. Additionally, the standard curves of all the metabolites were measured in the concentration range of 0.1, 1, 5, 10, and 50 mM. HPLC was performed using an Ultimate 3000 HPLC (Dionex) equipped with an RI-101 refractive index detector (Shodex, Kawasaki, Japan), an autosampler and an ion-exclusion Aminex HPX-87H column (7.8 mm × 300 mm) with a guard column (Bio-Rad, Hercules, CA). As the mobile phase 5 mM H_2_SO_4_ was used at a flow rate of 0.6 ml/min, the column was kept at 40°C. The total run time was 30 min and the injection volume was 10 μl.

### 2.3 Transmission Electron Microscopy (TEM)

*L. monocytogenes* EGDe cultures were grown anaerobically at 30°C rhamnose *pdu*-induced and rhamnose *pdu* non-induced condition. Samples were collected at 48 h of incubation. About 10 μg dry cells were fixed for 2 h in 2.5% (v/v) glutaraldehyde in 0.1 M sodium cacodylate buffer (pH 7.2). After rinsing in the same buffer, a post-fixation was done in 1% (w/v) OsO_4_ for 1 h at room temperature. The samples were dehydrated by ethanol and were then embedded in resin (Spurr HM20) 8 h at 70°C. Thin sections (<100 nm) of polymerized resin samples were obtained with microtomes. After staining with 2% (w/v) aqueous uranyl acetate, the samples were analyzed with a Jeol 1400 plus TEM with 120 kV setting [8, 9].

### 2.4 Proteomics

*L. monocytogenes* cultures were anaerobically grown at 30°C in rhamnose *pdu*-induced and rhamnose *pdu* non-induced condition. Samples were collected at 48 h of incubation and then washed twice with 100 mM Tris (pH 8). About 10 mg wet weight cells in 100 μl 100 mM Tris was sonicated for 30 s twice to lyse the cells. Samples were prepared according to the filter assisted sample preparation protocol (FASP) with the following steps: reduction with 15 mM dithiothreitol, alkylation with 20 mM acrylamide, and digestion with sequencing grade trypsin overnight [22]. Each prepared peptide sample was analyzed by injecting (18 μl) into a nanoLC-MS/MS (Thermo nLC1000 connected to a LTQ-Orbitrap XL) as described previously [8, 9]. LCMS data with all MS/MS spectra were analyzed with the MaxQuant quantitative proteomics software package as described before [8, 9, 23]. A protein database with the protein sequences of *L. monocytogenes* EGDe (ID: UP000000817) was downloaded from UniProt. Filtering and further bioinformatics and statistical analysis of the MaxQuant ProteinGroups file were performed with Perseus [24]. Reverse hits and contaminants were filtered out. Protein groups were filtered to contain minimally two peptides for protein identification of which at least one is unique and at least one is unmodified. The volcano plot was prepared based on the Student’s t-test difference of Pdu-induced/non-induced control.

### 2.5 Bioinformatics and Statistical Analysis

Pathview R package [25] to visualize the proteomics data: The UniProt protein IDs from Supplementary Table 1 were collected and retrieved to Entre IDs. A list of Entrez IDs, protein expression indicated by LFQ intensity (Supplementary Table 2) was mapped to the *L. monocytogenes* EGDe KEGG pathway database using the tool Pathview (R version 3.2.1). The box represent genes and the different color indicates level of expression with default setting.

Statistical analyses were performed in Prism 8.0.1 for Windows (GraphPad Software). As indicated in the figure legend, Statistical significances are shown in ***, P<0.001; *, P<0.05; ns, P>0.05 with Holm-Sidak T-test.

## 3. Results

### 3.1 Activation of *pdu* stimulates anaerobic growth of *L. monocytogenes* EGDe on rhamnose

We first examined if rhamnose can function as a sole carbon source to support anaerobic growth of *L. monocytogenes* EGDe in MWB defined medium without and with added vitamin B12 (cobalamin) (Figure 1). In MWB defined medium supplied with 20mM rhamnose OD_600_ reaches a maximum of about 0.37 after 48 h, while in MWB supplied with 20mM rhamnose and 20nM B12 OD_600_ continues to increase after 48 h, reaching a significant higher OD_600_ of 0.51 at 72 h. Enhanced growth on MWB supplied with rhamnose and B12 compared to MWB plus rhamnose, is also evident from plate counts which increase from 6.5 to 8.2 log10 CFU/ml and from 6.5 to 7.2 log10 CFU/ml, respectively (figure 1B). There is no significant difference in growth performance of *L. monocytogenes* EGDe on MWB supplied with 20mM glucose and MWB supplied with 20mM glucose and 20 nM B12, and at 48 h final levels of 8.8 log10 CFU/ml were reached (Supplementary Figure 1). These results suggest that in B12 stimulated anaerobic growth of *L. monocytogenes* EGDe on MWB medium with rhamnose as sole carbon source is linked to activation of *pdu*.

**Figure 1.**
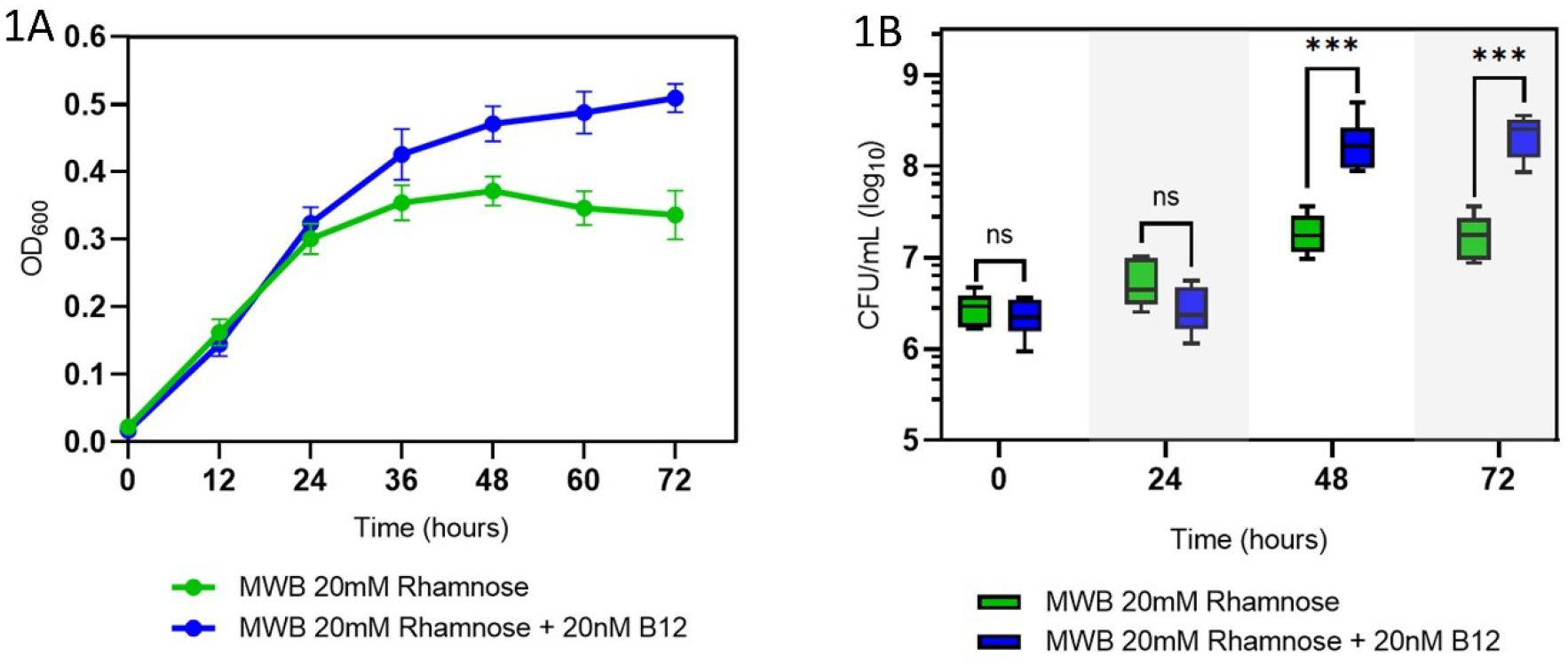
Impact of L-rhamnose and vitamin B12 on anaerobic growth of *L. monocytogenes* EGDe. (1A) OD_600_ growth curves in MWB defined medium with 20 mM L-rhamnose as sole carbon source (green symbols) and MWB with 20 mM Rhamnose and 20n M B12 (blue symbols). (1B) CFUs during growth on MWB 20mM Rhamnose (green symbols) and on MWB 20 mM Rhamnose + 20 nM B12 (blue symbols). Results from three independent experiments with three technical repeats are expressed as mean and standard errors. Statistical significance is indicated (***, P<0.001; ns, P>0.05 Holm-Sidak T-test).

### 3.2 Activation of *pdu* supports 1,2-propanediol degradation and stimulates rhamnose metabolism

To confirm possible activation of *pdu*, metabolic analysis via HPLC was conducted to quantify substrate consumption and product formation following anaerobic growth of *L. monocytogenes* EGDe on MWB plus 20mM rhamnose and MWB plus 20mM rhamnose and 20nM B12. As shown in Figure 2A, at 72 h, the initial 20mM rhamnose is completely consumed in *pdu* induced condition, whereas 3.5 mM of rhamnose is retained in *pdu* non-induced condition. Additional end product analysis at 72 h shows accumulation of approximately 6.7 mM of 1,2-propanediol in *pdu* non-induced condition and nearly zero production of propionate and 1-propanol. In *pdu* induced condition a significant lower amount of 1,2-propanediol is found, about 1.4 mM, and higher levels of approximately 3.4 mM propionate and 3.6 mM 1-propanol are produced at 72 h, in line with the expected 1:1 molar stoichiometry of *L. monocytogenes* BMC-dependent pdu [9]. Enhanced rhamnose metabolism in *pdu*-induced cells is also evident from production of acetate and lactate. At 72h, 4.1 mM acetate and 2.3 lactate are produced in *pdu* non-induced condition while 7.6 mM acetate and 5.1 mM lactate are produced in *pdu* induced condition.

**Figure 2.**
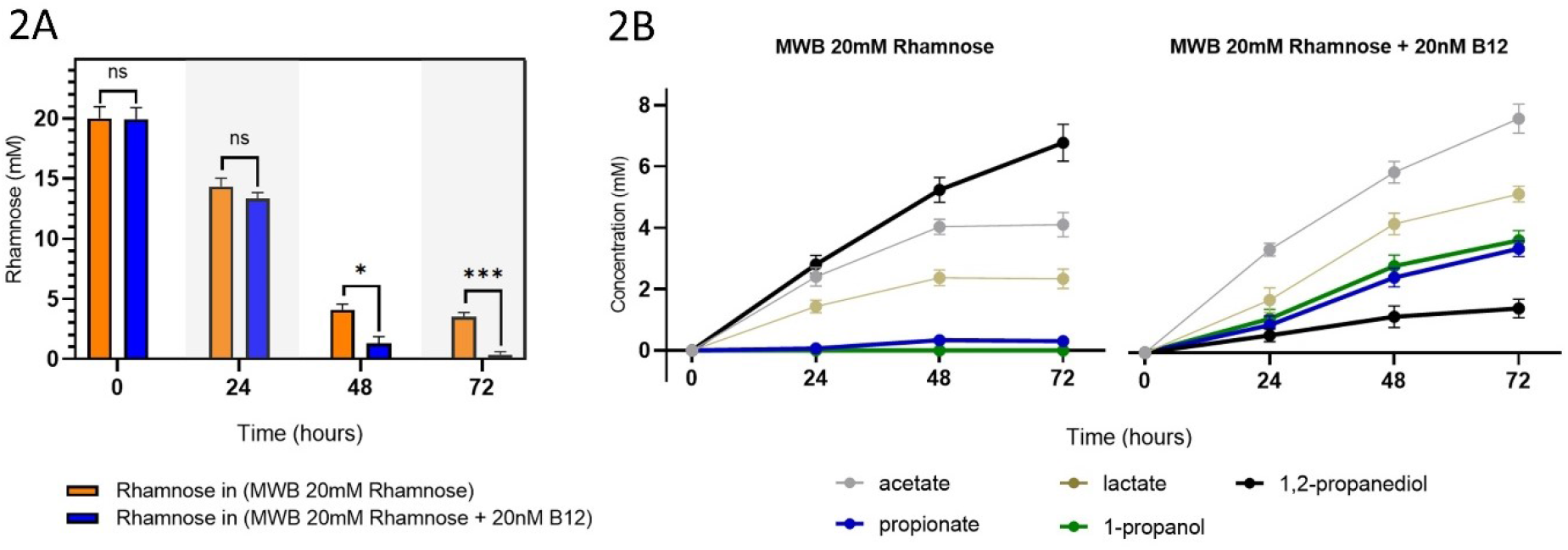
Impact of vitamin B12 on Rhamnose metabolism of anaerobically grown *L. monocytogenes* EGDe. (2A) Utilization of Rhamnose by *L. monocytogenes* EGDe anaerobically grown in MWB plus 20 mM Rhamnose (orange bars) and MWB plus 20mM Rhamnose and 20 nM B12 (blue bars). (2B) Metabolites from Rhamnose metabolism of *L. monocytogenes* EGDe anaerobically grown in MWB plus 20mM Rhamnose (left) and MWB plus 20 mM Rhamnose and 20 nM B12 (right). Results from three independent experiments are expressed as mean and standard errors. Statistical significance is indicated (***, P<0.001; *, P<0.05; ns, P>0.05 Holm-Sidak T-test).

### 3.3 Visualization of BMCs and expression analysis of BMC shell proteins

To answer the question if BMCs are formed to support the utilization of rhamnose-derived 1,2-propanediol, Transmission Electron Microscopy (TEM) was performed to observe BMCs structures, and proteomics was applied to measure the expression of BMC shell proteins (Figure 3A). The *pdu* induced cells clearly contain BMC-like structures with an approximate diameter of 50–80 nm, which are absent in *pdu* non-induced cells. Notably, the identified structures strongly resemble TEM pictures of previously reported pdu BMCs in *L. monocytogenes* [8, 9] and in *S. enterica* and *E. coli* [13, 26]. Compared to *pdu* non-induced cells, *pdu* induced cells show significant upregulation of 21 measurable Pdu proteins (Figure 3B), including seven proteins annotated as BMCs shell proteins, PduTUABKJN. Notably, *pdu*-induced and *pdu* non-induced rhamnose grown cells show similar expression of proteins in the rhamnose metabolism cluster (*lmo2850, rhaA, rhaB* and *rhaM*) (Figure 3B), which indicates that the activation of pdu BMC does not affect the expression of these enzymes.

**Figure 3.**
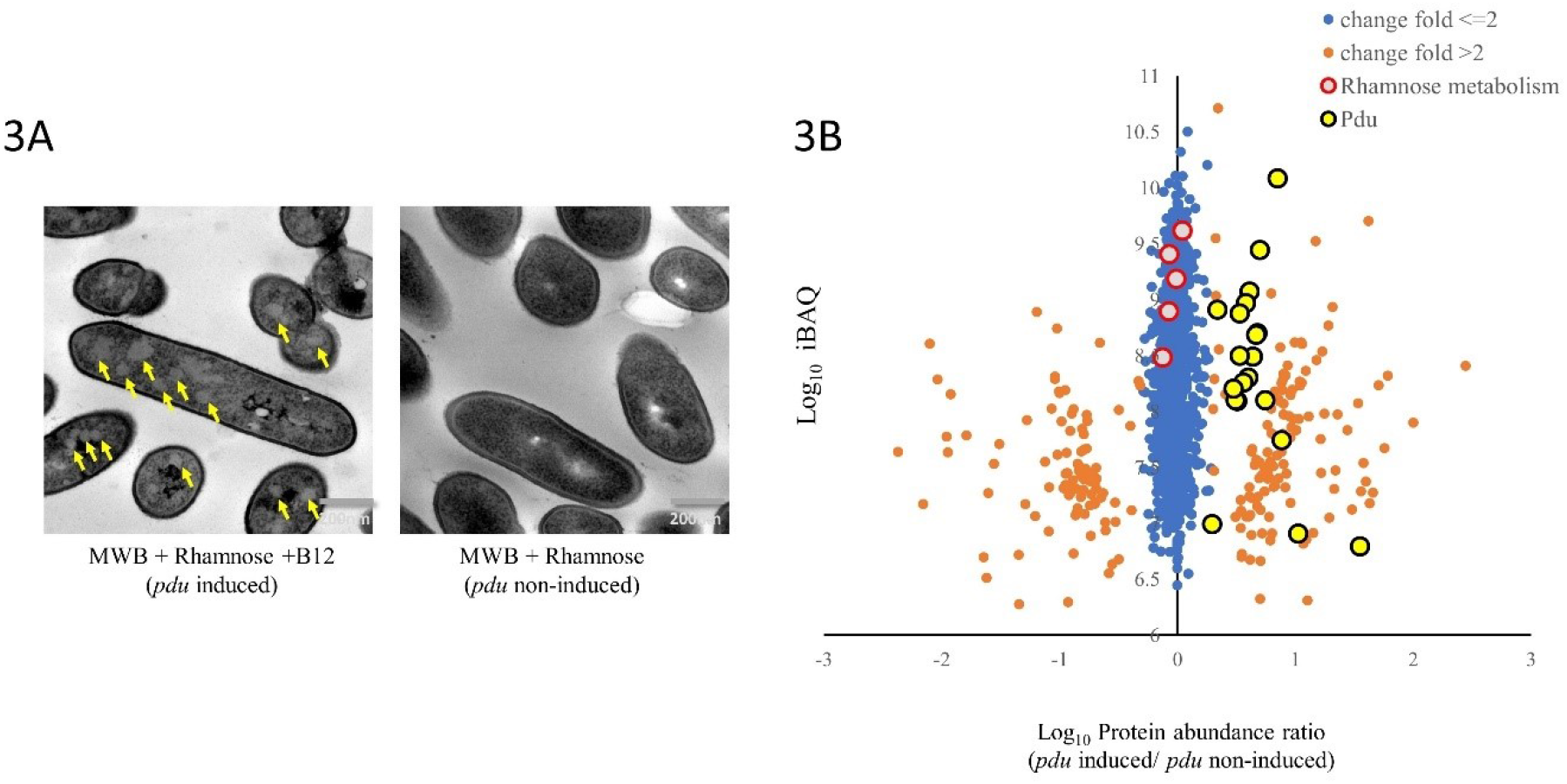
TEM visualization of BMCs and proteomics analysis of *pdu* induced cells (MWB plus 20 mM Rhamnose and B12) compared to *pdu* non-induced cells (MWB plus 20 mM Rhamnose). (3A) TEM visualization of BMCs in cells grown on MWB plus 20 mM Rhamnose and B12 (left; yellow arrows point to BMCs) and cells grown on MWB with 20mM Rhamnose (right). (3B) Proteomic volcano plot of MWB plus 20mM Rhamnose and B12 compared to MWB plus 20 mM Rhamnose grown cells. Fold change ≤ 2 in blue, fold change > 2 in orange, proteins in the Pdu cluster are black encircled in yellow, and proteins in the Rhamnose cluster are red-encircled in grey. More details in text and in Supplementary Table 1.

### 3.4 Proteomics-based pathway visualization of propanoate metabolism and vitamin B12 metabolism

To visualize the metabolism from 1,2-propanediol to propanoate (propionate) and 1-propanol, the identified proteins and expression levels presented in Supplementary Table 1, are mapped to propanoate metabolic pathways of *L. monocytogenes* EGDe. As shown in Figure 4A, the enzymes involved in degradation of rhamnose-derived 1,2-propanediol into propanoate (propionate) and 1-propanol are all significantly upregulated in *pdu* induced condition compared to *pdu* non-induced condition. The propanediol dehydratase (EC 4.2.1.28) is an enzyme with three subunits encoded by *pduC, pduD* and *pduE*, which converts 1,2-propanediol into propanal (propionaldehyde). Propionaldehyde is metabolized to 1-propanol by propanol dehydrogenase PduQ and propanol-CoA by propionaldehyde dehydrogenase PduP (EC 1.2.1.87). Propanol-CoA is converted to propanoyl-phosphate by phosphate propanoyltransferase PduL (EC 2.3.1.222), with propanoyl-phosphate subsequently converted to propanoate by propionate kinase PduW (EC 2.7.2.1). We found that the vitamin B12 biosynthesis pathway that is grouped in porphyrin and chlorophyll metabolism, is significantly downregulated in *pdu* induced condition compared to *pdu* non-induced condition (Figure 4B), which suggests that supplementation of 20nM B12 represses the expression of proteins required for B12 biosynthesis. This also includes the three enzymes mediating the final steps in B12 biosynthesis, CobU, CobS and CobC, encoded by the respective genes located in the *pdu* cluster (Figure 4B) [8, 27–29]. Apparently, B12 accumulation from the medium supports activation of *pdu* BMCs, whereas despite expression of B12 biosynthesis enzymes, production of B12 and levels reached are not sufficient to induce *pdu* in *L. monocytogenes* EGDe grown in MWB without added B12.

**Figure 4.**
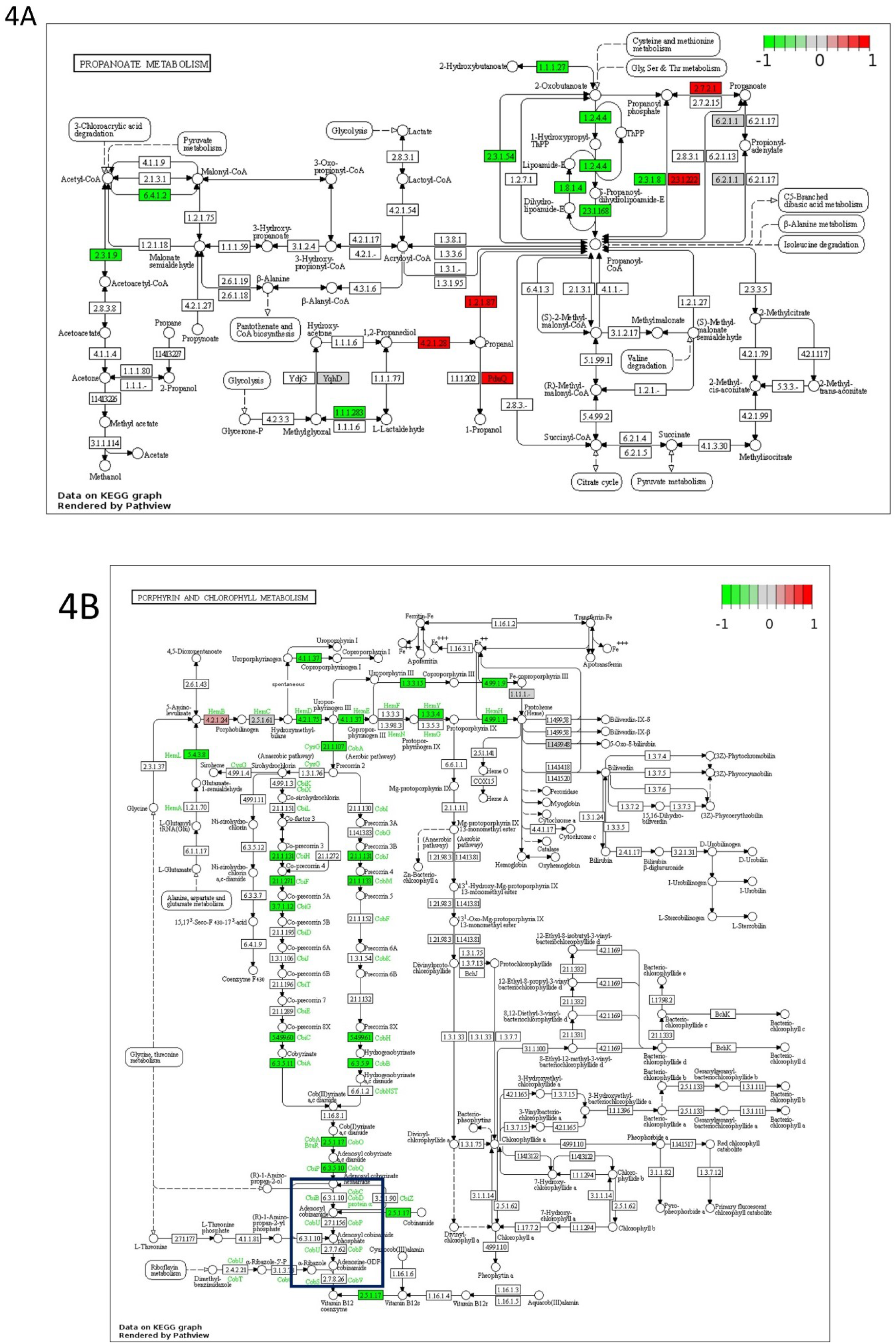
Proteomics-based pathway visualization of propanoate metabolism (4A) and porphyrin and chlorophyll metabolism (4B) in *pdu* induced compared to *pdu* non-induced *L. monocytogenes* EGDe via Pathview. Rectangle boxes represent enzymes with the relative expression indicated based on proteomics data. Key metabolites are named and positions in the pathways indicated by circles. In (4B), the blue box highlights B12 reactions that encoded by *pdu* cluster, more details in text and in Supplementary Table 4.

## 4. Discussion

The presented model of 1,2-propanediol BMCs in rhamnose metabolism is based on growth phenotypes, metabolic analysis, proteomics, TEM visualization and our understanding of 1,2-propanediol BMCs in anaerobic growth of *L. monocytogenes* EGDe. As illustrated in Figure 5, the rhamnose catabolism gene cluster (*rha*) in *L. monocytogenes* EGDe is composed of *lmo2846-lmo2851* [30]. *lmo2850* encodes a secondary transporter which has high similarity with L-rhamnose permease RhaT in *E. coli* [31–33], and is conceivably acting as the transporter of α-L-rhamnose. L-rhamnose mutarotase RhaM mediates the conversion of α-L-rhamnose into β-L-rhamnose (also called L-rhamnopyranose) [30, 34]. β-L-rhamnose is converted to L-rhamnulose by L-rhamnose isomerase RhaA [30, 35]. L-rhamnose is then phosphorylated to L-rhamnulose 1-phosphate by rhamnulokinase RhaB with one ATP consumption [30, 35]. L-rhamnulose 1-phosphate is split into (S)-lactaldehyde and dihydroxyacetone phosphate (DHAP) by rhamnulose-1-phosphate aldolase RhaD [30, 35]. DHAP can be metabolized to glyceraldehyde 3-phosphate via Triosephosphate isomerase 1 TpiA1 and via the glycolytic pathway [14, 36] and the GABA (γ-aminobutyric acid) shunt in the incomplete TCA cycle in *L. monocytogenes* [37], to the end products acetate and lactate, as confirmed in our metabolic analysis. The observed production of 1,2-propanediol in *pdu* non-induced conditions confirms the predicted anaerobic conversion of lactaldehyde to 1,2-propanediol in *L. monocytogenes* EGDe. The activity of lactaldehyde reductase has not been described in *L. monocytogenes* [38], but protein similarity alignment with lactaldehyde reductase FucO of *Escherichia coli* [38], suggests four putative candidates annotated as alcohol dehydrogenase in *L. monocytogenes* EGDe including lmo1166, lmo1171, lmo1634 and lmo1737, detected in the proteomes of both *pdu* non-induced and *pdu*-induced cells (for details see Supplementary File 1). Since the discovery of the role of *pdu* BMCs dehydratase in rhamnose (and fucose) utilization, two pathway scenarios have been proposed, one with and one without lactaldehyde reductase encapsulated inside BMCs [30, 34]. In line with previously reported comparative genomic analysis [30, 34], our data now provide evidence for the latter model to be active in *L. monocytogenes* since rhamnose is converted via lactaldehyde to 1,2-propanediol in absence of BMCs in *pdu* non-induced condition, while with added B12 metabolism of 1,2-propanediol proceeds via *pdu* BMCs.

**Figure 5.**
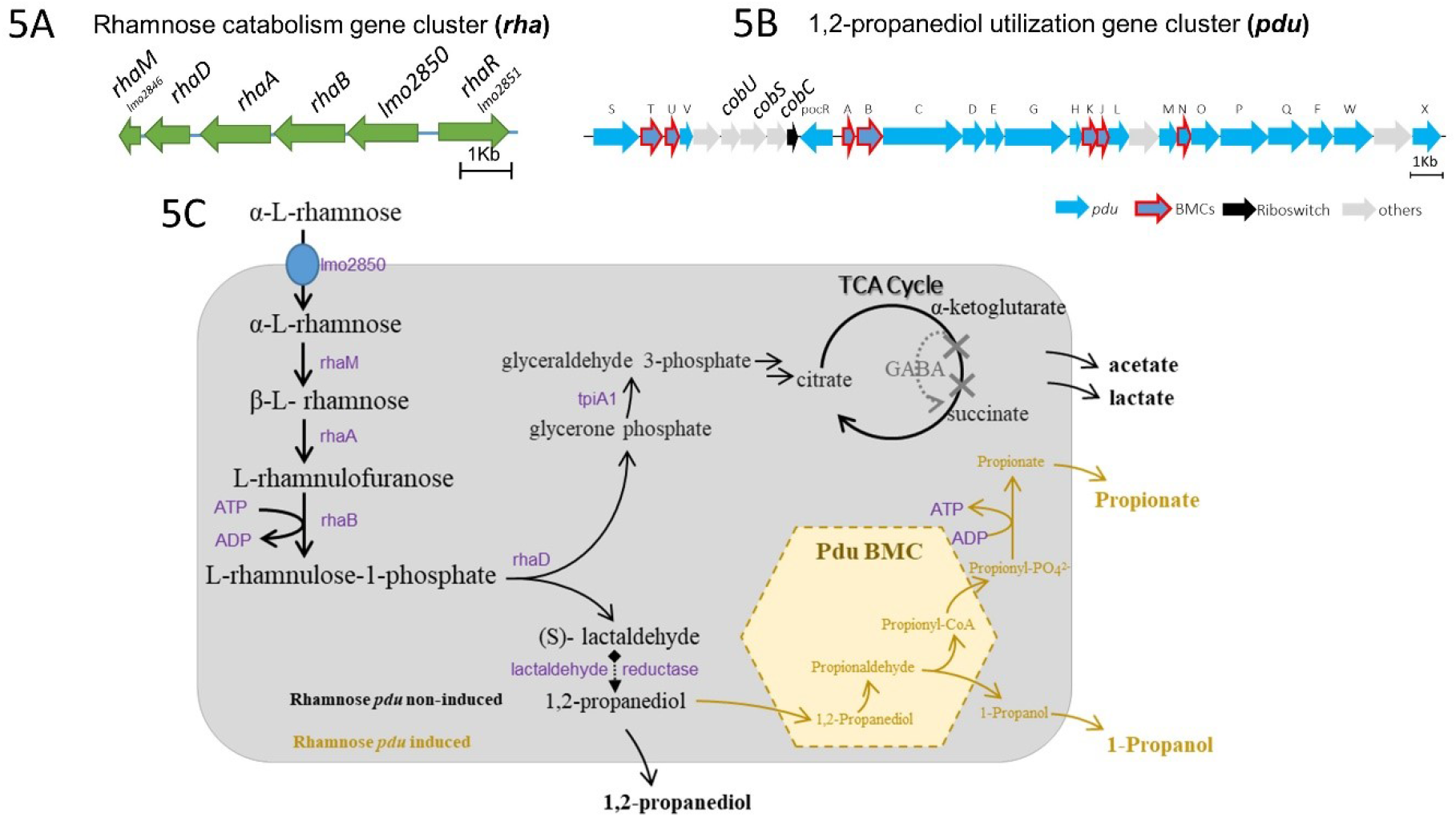
Overview of rhamnose metabolism with or without 1,2-propanediol BMCs in *L. monocytogenes*. (5A) Rhamnose catabolism gene cluster, *rha* (5B) 1,2-propanediol utilization gene cluster, *pdu*. Details for (5A) and (5B) are in Supplementary Table 4. (5C) The Proposed rhamnose metabolism model based on this study. Arrows represent reactions and enzymes and compounds indicated in black represent rhamnose metabolism without BMCs, and 1,2-propanediol BMC reactions activated by B12 and compounds involved are shown in yellow. For details see corresponding sections in results and discussion.

The activation of *pdu* BMCs enhances anaerobic rhamnose metabolism in *L. monocytogenes* and generates additional energy via the ATP producing propionate branch in *pdu*, and via enhanced flux into the glycolytic pathway resulting in a significant stimulation of growth. At 72 h, 20 mM rhamnose is metabolized into 7.6 mM acetate, 5.1 mM lactate, 1.4mM 1,2-propanediol, 3.4 mM propionate and 3.6 mM 1-propanol in *pdu* induced condition, while 16.5 mM rhamnose is metabolized into 4.1 mM acetate, 2.3 mM lactate, 6.7 mM 1,2-propanediol in *pdu* non-induced condition. Theoretical ATP yield from rhamnose conversion to lactate, acetate and propionate includes production of 1.5 ATP per 1 lactate, 2.5 ATP per 1 acetate and 0.5 ATP per 1 propionate produced (for details of reactions see Supplementary Table 3). Based on concentrations of end products at 72h, *pdu* induced cells theoretically generate 1.425 ATP per 1 rhamnose while *pdu* non-induced cells generate 0.830 ATP per 1 rhamnose (for details of calculations see Supplementary Table 3). The improved energy gain of *L. monocytogenes* EGDe from anaerobic rhamnose metabolism with the activation of 1,2-propanediol BMCs, offers an explanation for the 10-fold higher number of CFUs reached (8.2 log10 CFU/ml) compared to non-pdu induced conditions (7.2 log10/ml).

Our data provide evidence for another extension of the BMC dependent metabolic repertoire of *L. monocytogenes* in anaerobic conditions, that now includes BMC-dependent ethanolamine utilization (eut) [9], BMC *pdu* [8], and BMC *pdu*-stimulated rhamnose metabolism. The indicated substrates can be found in a wide range of environments including foods and human gastrointestinal tract. In the latter case, competitive fitness of *L. monocytogenes* in the human intestine may be enhanced by supply of these compounds by enzymatic activity including release of ethanolamine following membrane phospholipid degradation and release of rhamnose following mucus glycan hydrolysis activity of for example *Bacteriodes spp*, and propanediol as a fermentation product [15]. Notably, despite the presence of a complete vitamin B12 synthesis cluster, we found that *eut* [9], *pdu* [8], and *pdu*-stimulated rhamnose utilization in *L. monocytogenes* in the current study, requires supplementation of B12 to the medium. This points to an important role of B12 in activation of *L. monocytogenes* BMC-mediated metabolic pathways containing B12-dependent signature aldehyde reductases. Vitamin B12 can be found in foods including meat and dairy products [28, 39] and is also found in human intestine, where part of the B12 is derived from gut microbiota that have the capacity to produce B12 [12, 28]. The fact that we observed in the current study induction of the B12 synthesis pathway in cells grown in MWB plus rhamnose but no activation of B12-dependent *pdu*, while activation was found with B12 added to the medium, points to an intricate regulation of the B12 synthesis pathway and its connection to BMCs activation. In addition to earlier studies on transcriptional and translational control of BMC *eut* and *pdu* in *L. monocytogenes* [12, 28, 40], studies are required to assess for example impact of extracellular and intracellular B12 concentrations on BMC pathway activation and their role in *L. monocytogenes* ecophysiology and virulence.

---

**Supplementary Figure 1.**
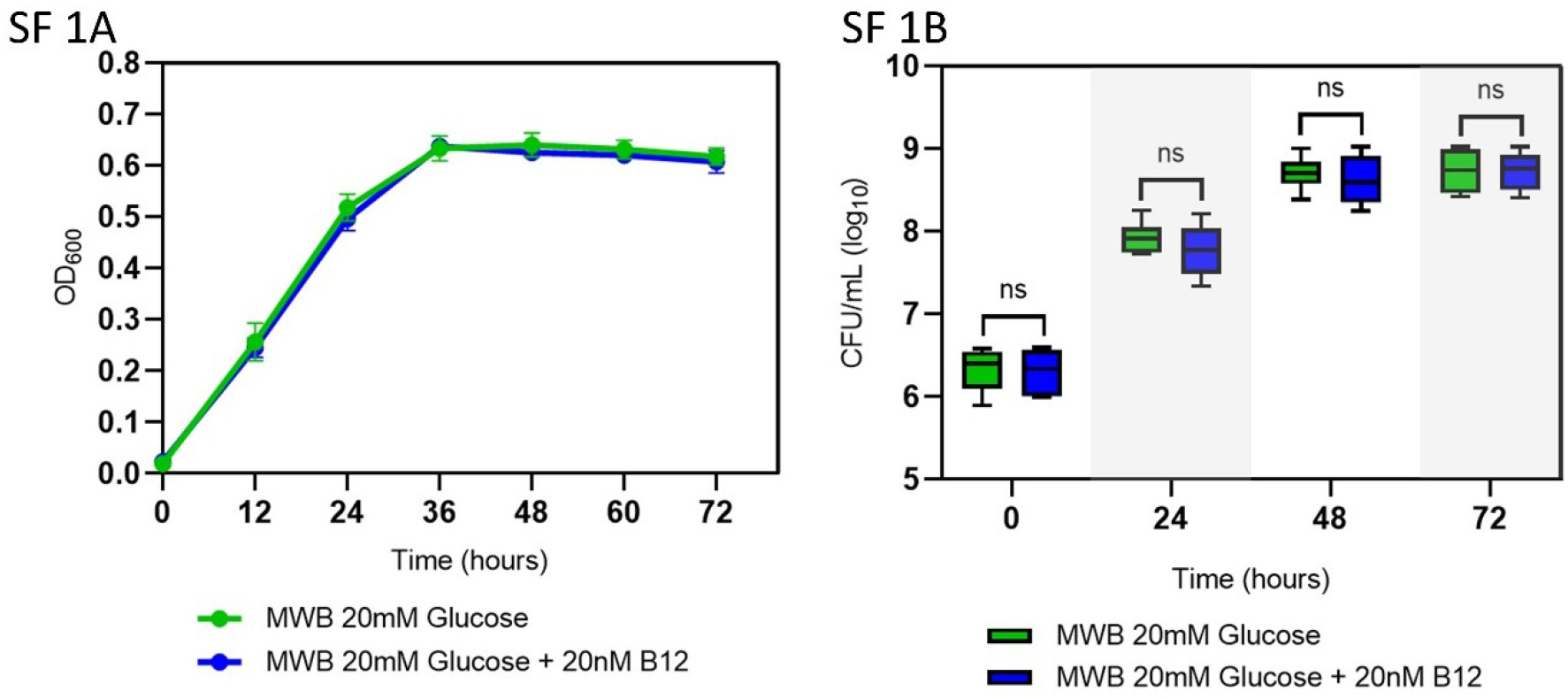
Anaerobic growth of *L. monocytogenes* EGDe on MWB plus glucose (green symbols) and MWB plus 20 mM glucose and 20 nM B12 (blue symbols). (SF 1A) OD600 growth curves, (SF 1B) CFUs determined at indicated time points during growth on MWB plus 20 mM glucose (green bars) and MWB plus 20 mM glucose and 20 nM B12 (blue bars). Results from three independent experiments with three technical repeats are expressed as mean and standard errors. Statistical significance is indicated (ns, P>0.05 Holm-Sidak T-test)

## 6. Supplementary Materials

Supplementary Table 1. Protein profiling of *pdu*-induced compared with non-induced *L. monocytogenes* EGDe in MWB medium with Rhamnose.

Supplementary Table 2. Input to Pathview with Entrez IDs and protein expression indicated by LFQ intensity.

Supplementary Table 3. Reaction list of Rhamnose metabolism and theoretical ATP yield from rhamnose conversion to lactate, acetate and propionate.

Supplementary Table 4. Annotation and Proteins IDs of *rha* and *pdu* cluster

Supplementary File 1. Protein similarity alignment with lactaldehyde reductase FucO of *Escherichia coli* against *L. monocytogenes* EGDe

